# Biodegradable harmonophores for targeted high-resolution *in vivo* tumor imaging

**DOI:** 10.1101/694760

**Authors:** Ali Yasin Sonay, Sine Yaganoglu, Martina Konantz, Claire Teulon, Sandro Sieber, Shuai Jiang, Shahed Behzadi, Daniel Crespy, Katharina Landfester, Sylvie Roke, Claudia Langerke, Periklis Pantazis

**Affiliations:** Department of Biosystems Science and Engineering (D-BSSE), Eidgenössische Technische Hochschule (ETH) Zurich, 4058 Basel, Switzerland; Department of Biomedicine, University Hospital Basel and University of Basel, Basel, Switzerland; Laboratory for Fundamental BioPhotonics, Institute of Bioengineering, School of Engineering, École Polytechnique Fédérale de Lausanne, CH-1015 Lausanne, Switzerland; Division of Pharmaceutical Technology, Department of Pharmaceutical Sciences, University of Basel, Basel, Switzerland; Max Planck Institute for Polymer Research, 55128 Mainz, Germany; Department of Materials Science and Engineering, School of Molecular Science and Engineering, Vidyasirimedhi Institute of Science and Technology (VISTEC), Rayong 21210, Thailand; Division of Hematology, University Hospital Basel, Basel, Switzerland; Department of Bioengineering, Imperial College London, South Kensington Campus, London SW7 2AZ, UK

**Author notes:** Correspondence to: Periklis Pantazis.

## Abstract

Optical imaging probes have played a major role in detecting and monitoring of a variety of diseases^1^. In particular, nonlinear optical imaging probes, such as second harmonic generating (SHG) nanoprobes, hold great promise as clinical contrast agents, as they can be imaged with little background signal and unmatched long-term photostability^2^. As their chemical composition often includes transition metals, the use of inorganic SHG nanoprobes can raise long-term health concerns. Ideally, contrast agents for biomedical applications should be degraded *in vivo* without any long-term toxicological consequences to the organism. Here, we developed biodegradable harmonophores (bioharmonophores) that consist of polymer-encapsulated, self-assembling peptides that generate a strong SHG signal. When functionalized with tumor cell surface markers, these reporters can target single cancer cells with high detection sensitivity in zebrafish embryos *in vivo*. Thus, bioharmonophores will enable an innovative approach to cancer treatment using targeted high-resolution optical imaging for diagnostics and therapy.

## Main Text

Clinical and preclinical imaging holds great potential in mapping disease progression and can provide diagnostic information that may guide the choice of treatment strategies for disease^3, 4^. Optical techniques using bioluminescent and fluorescent probes have emerged as promising modalities for molecular imaging in disease and therapy due to their ease of use and improved cellular resolution, capable of distinguishing boundaries between malignant and normal tissue^5^. A key challenge for optical imaging probes and instrumentation, particularly those aimed at eventual clinical applications, is to overcome the limited depth penetration of excitation light, which often result in a low signal-to-noise ratio (SNR)^6^. The relatively poor photostability of most imaging probes pose another challenge to provide reliable and sensitive imaging of tumors.

Previously, we introduced inorganic second harmonic generating (SHG) nanocrystals, SHG nanoprobes^2^, as a new class of imaging probes that can be used for *in vivo* imaging. Given that SHG imaging employs near infrared (NIR) incident light for contrast generation, SHG nanoprobes can be utilized for deep tissue imaging. Unlike commonly used fluorescent probes, SHG nanoprobes neither bleach nor blink, and their signal does not saturate with increasing illumination intensity, ensuring high probe sensitivity^7^. Since their signal profile is very narrow, they can be imaged with high SNR by excluding the broad emission of typical autofluorescence background^2^. Robust functionalization allows targeting to a wide variety of cells and proteins of interest^8^, allowing these imaging probes to be promising tools for both clinical and preclinical imaging applications^9^. Despite these advantages, the chemical structure of inorganic SHG nanoprobes makes them stable in the body, which may cause concerns for the long-term health of an organism that has been imaged with these reporters.

To create a foundation for safe SHG nanoprobe-based clinical imaging, we set out to generate a nanoprobe that consists of biodegradable materials, capable of generating sufficient SHG signal that can be detected with high SNR. Our efforts were guided by the observation that peptides with a variable number of amino acid units can self-assemble into large, solid nanostructures of different morphologies and symmetries^10^ (**Fig. 1a)**. It has been previously shown that such nanostructures can be ferroelectric and give nonlinear optical contrast such as SHG^11, 12^ (**Fig. 1c**).

**Figure 1.**
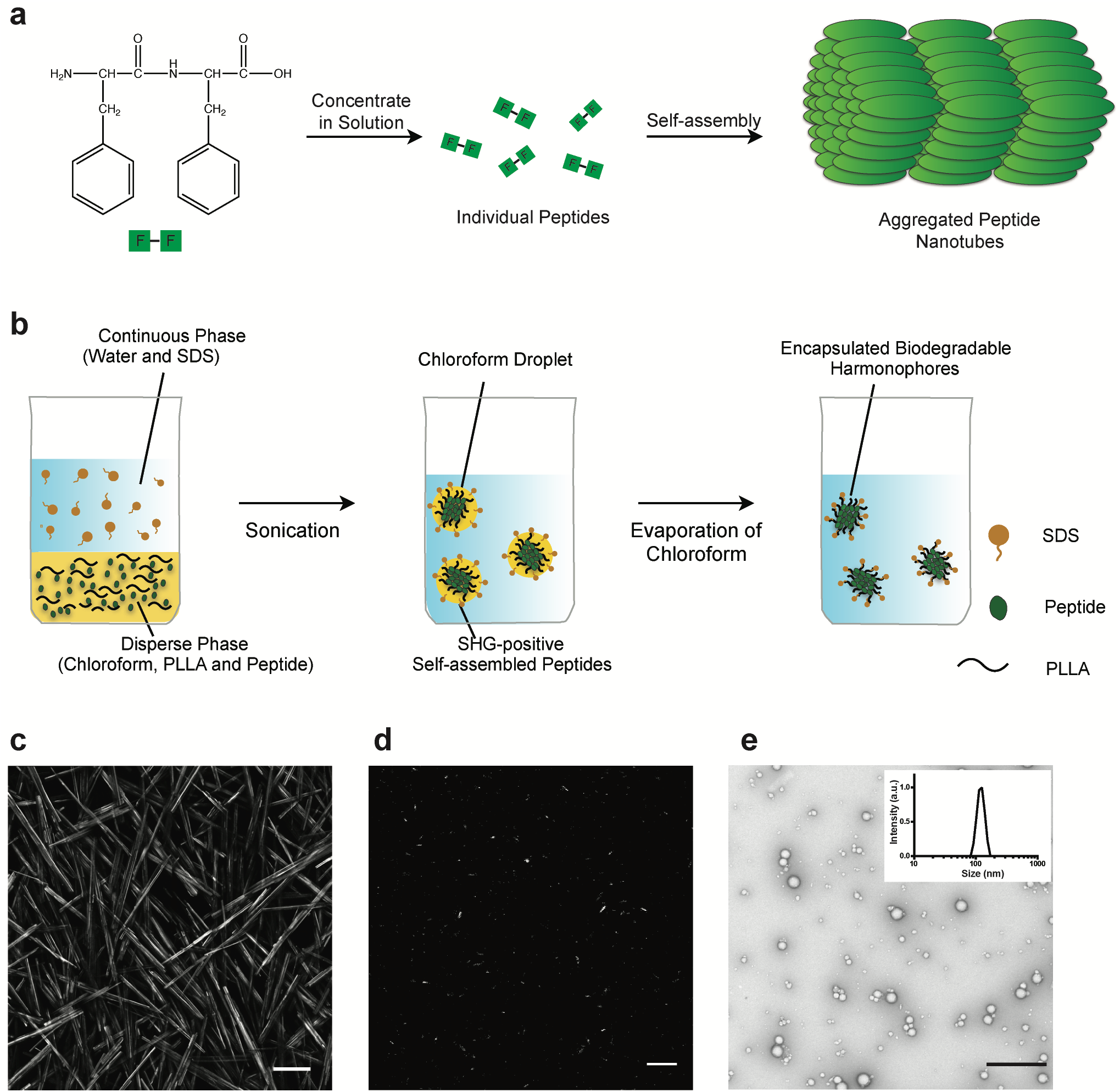
Synthesis and analysis of bioharmonophores. **a**, Schematic of the self-assembling reaction of diphenylalanine peptides (FF) into large-scale nanotube structures from a concentrated solution. **b**, Schematic of the emulsion-solvent evaporation method for the synthesis of bioharmonophores. Self-assembling peptides are dissolved in chloroform along with biodegradable poly(*L*-lactic acid) (PLLA) and emulsified with the surfactant sodium dodecyl sulfate (SDS) using sonication, followed by evaporation of chloroform. **c**, SHG signal from diphenylalanine peptide nanotubes aggregated on top of the imaging chamber. Peptide nanotubes were illuminated with a 850 nm pulsed laser. Image composite of multiple stitched images. **d**, SHG signal from encapsulated triphenylalanine peptides (FFF) bioharmonophores immobilized in 1% low melting agarose illuminated with 850 nm pulsed laser. **e**, TEM image of synthesized FFF-based bioharmonophores showing uniform spherical nanoparticles. **Inset**: DLS data showing the size distribution of synthesized bioharmonophores. Scale bar, 100 μm (**c**); Scale bar, 10 μm (**d**); Scale bar, 500 nm (**e**).

To render these nanostructures suitable for biological applications, we evaluated methods for the encapsulation of self-assembling peptides in order to hinder their macroscopic aggregation by confining their self-assembly in nanodroplets without affecting their ability to generate a strong SHG signal, and to generate a nanoparticle that can be further functionalized without influencing the peptide assembly. To this end, we subjected several peptides that have been reported to self-assemble into complex nanostructures to the emulsion-solvent evaporation method^13^, a widely-used procedure for the fabrication of monolithic and core–shell nanoparticles (**Fig. 1b**, **see Methods**).

We identified three peptides with different self-assembling properties (pentaalanine^14^, trileucine^15^, and triphenylalanine^16^) that could generate detectable SHG signal when encapsulated in the biodegradable polymer (**Fig. 1d** and **Supplementary Fig. 1**). Transmission electron microscopy (TEM) analysis of the predominantly spherical nanoparticles, hereinafter referred to as bioharmonophores, revealed a diameter ranging from 50-150 nm, which was confirmed by dynamic light scattering (DLS) measurements (**Fig. 1e)**.

SHG signal from bioharmonophores can stem from i) the bulk of the self-assembling peptides that form noncentrosymmetric crystalline structures or ii) the surface of the bioharmonophores where there is no inversion symmetry. To ascertain that the SHG signal originates from the crystalline peptide core, we performed X-ray diffraction (XRD) analysis of bioharmonophores with different peptide contents. In all cases, the peptides showed a high degree of internal order with distinct diffraction patterns associated with their individual crystalline phases and self-assembling behavior (**Supplementary Fig. 2a-d**).

Because bioharmonophores based on triphenylalanine (FFF) peptides yielded the strongest SHG signal compared to pentaalanine and trileucine, we subjected these bioharmonophores to detailed optical characterizations. The SHG signal of FFF-based bioharmonophores was spectrally well-defined (**Fig. 2a**). Additionally, the SHG emission patterns of FFF-based bioharmonophores displayed a broad opening: one seemingly isotropic, and the other one displaying one lobe over 60° in the forward direction (**Fig. 2b**). These results indicate that bioharmonophores emit SHG signal in multiple directions (unlike the predominantly forward-directed SHG of large protein arrays)^7^, which allowed illumination and collection of SHG signal using the same microscope objective lens. Moreover, the presence of a single lobe demonstrates that the observed SHG signal originates from the bulk of the bioharmonophores and not from its surface, as described by Mie theory^17^.

**Figure 2.**
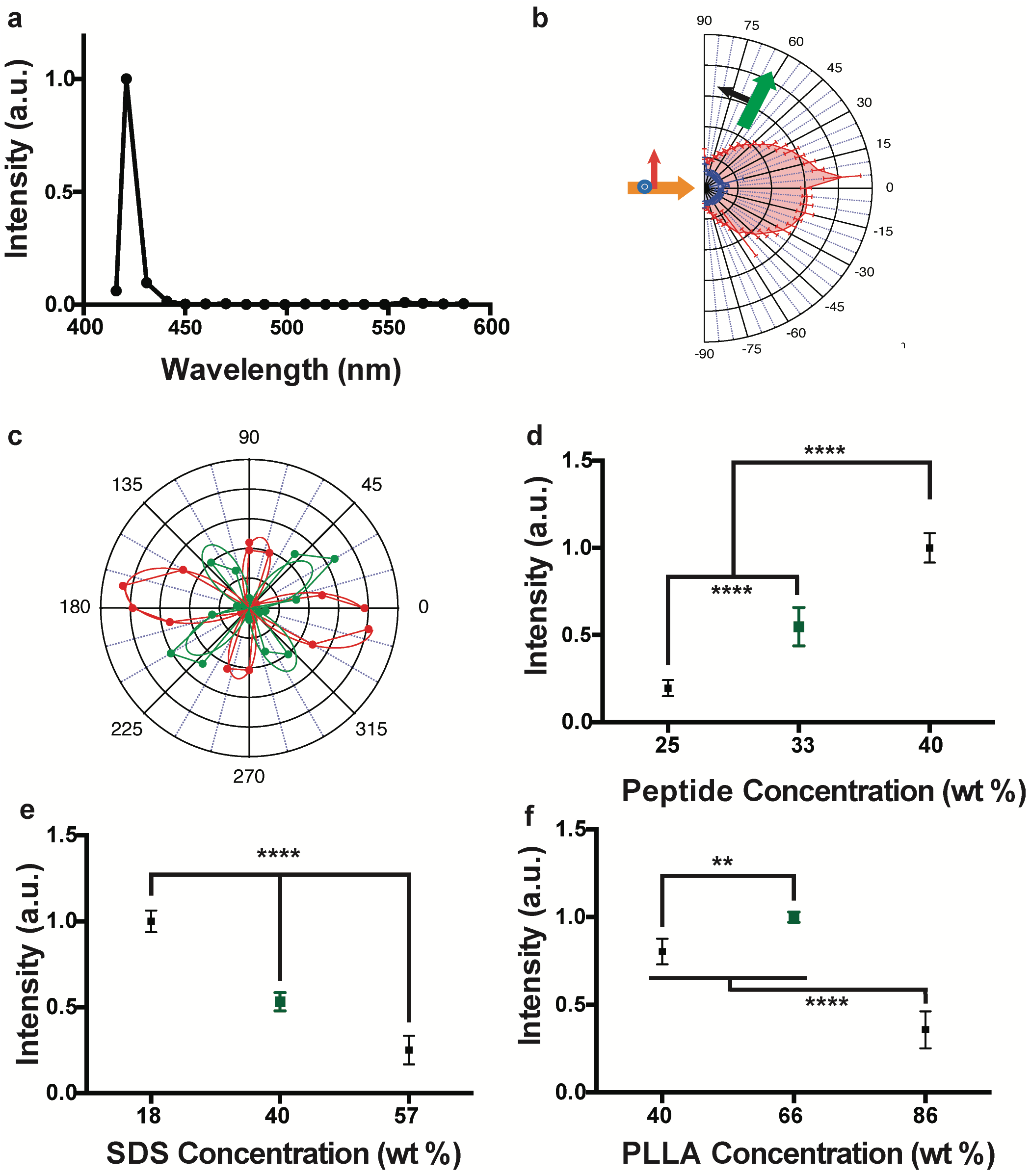
Optical characterization of bioharmonophores and analysis of parameters influencing harmonophore formation. **a**, Normalized SHG signal spectrum of FFF-based bioharmonophores (signal ranging from 400 to 600 nm) illuminated with 850 nm pulsed laser. The characteristic SHG peak is centered around 425 nm. **b**, SHG emission pattern of Triphenylalanine based bioharmonophores. Orange arrow indicates excitation beam direction. Green arrow shows SHG collection direction, which rotates between −90° and 90°. The detected polarization is in the beams plane (P, black arrow). Red pattern shows PPP polarization configuration (excitation and detection polarizations in the beams plane), and blue pattern shows PSS (excitation with a perpendicular polarization). **c**, SHG intensity vs. incident polarization angle for a bioharmonophore, highlighted by the solid white circle in **Supplementary Fig. 3**. Red color shows detection along the X axis while green color shows detection along the Y axis. The experimental curve is a dotted line, the corresponding fitted curve, assuming C_2_ symmetry, is a solid line. **d**, Influence of using different amounts of FFF peptide during bioharmonophore production on the SHG signal intensity. The optimal condition (33 wt%) is marked in green. The use of higher FFF peptide amount leads to aggregates (n=5). **e**, Influence of SDS concentration (wt% of disperse phase) on SHG intensity of generated bioharmonophores. The optimal condition (40 wt% SDS) with high bioharmonophore stability and less aggregation is marked in green (n=5). **f**, Influence of using different amounts of PLLA during bioharmonophore production on the SHG intensity of the generated bioharmonophores. The optimal condition (66 wt% PLLA) is marked in green (n=5). Mean ± s.d. ****, P < 0.0001, **, P < 0.005, *, P < 0.05 (Ordinary one-way ANOVA with Tukey’s multiple comparisons).

Because SHG involves only virtual energy transitions, bioharmonophores did not display blinking, remained stable over extended periods of illumination, and their SHG signal intensity rose quadratically when the laser intensity shone on them was linearly increased (**Supplementary Fig. 3a,b**). The measured polarimetric diagrams (**Fig 2c and Supplementary Fig. 3c-f**) were consistent with the hypothesis that bioharmonophores have a self-assembling peptide core with a monoclinic (C2) symmetry. Indeed, the experimental curves were well fitted with the analytical expression calculated for this symmetry (see **Supplementary Note 1** – Optical Characterization of Bioharmonophores). Taken together, bioharmonophores have the same photophysical advantages for biomedical imaging applications that have been previously described for inorganic SHG nanoprobes^2^.

To gain insight into the parameters influencing the bioharmonophore stability and signal intensity, we tested several reaction conditions to generate bioharmonophores. Given that the SHG signal originating from bioharmonophores is dependent on the amount of encapsulated peptide, we first tested whether varying the FFF peptide concentration during production would improve the SHG signal intensity of generated bioharmonophores (**Fig. 2d and Supplementary Fig. 4**). We found that an amount of 15 mg FFF (i.e. 33wt%) peptide provided an optimal combination of intense SHG signal and bioharmonophore stability. Interestingly, while an FFF peptide amount of 20 mg (40wt%) increased the overall SHG signal, it also led to bioharmonophore aggregation and decreased colloidal stability. Conversely, 10 mg (25wt%) FFF peptide generated little SHG signal.

Because surfactant concentration plays a crucial role in emulsification of chloroform droplets^13^, we reasoned that altering the surfactant concentration during the preparation of bioharmonophores would have a profound effect on their stability and signal strength (**Fig. 2e and Supplementary Fig. 5)**. Bioharmonophores emulsified in an aqueous solution with 0.3% sodium dodecyl sulfate (SDS) (i.e. 40wt% of dispersed phase) yielded stable bioharmonophores with intense SHG signal, whereas compositions employing 0.1% SDS (18wt%) yielded aggregated nanoparticles. Increasing the SDS concentration to 0.6% (57wt%) diminished the SHG signal intensity, suggesting that the bioharmonophore size and hence the number of enclosed peptide molecules within each bioharmonophore is influenced by the concentration of surfactant.

Finally, we varied the polymer quantity that encapsulates and shields peptides from environmental changes, and assessed its role in both SHG signal intensity and nanoparticle morphology (**Fig. 2f and Supplementary Fig. 6**). We identified that an amount of 30 mg of poly(*L*-lactic acid) (PLLA) (66wt%) resulted in an optimal combination of intense SHG signal and bioharmonophore stability. Lower polymer amount of 10 mg (40 wt%) yielded weaker SHG signal, whereas higher polymer amount (90 mg, 86 wt%) led to elongated bioharmonophore morphologies. Taken together, we identified optimal experimental conditions to generate bioharmonophores providing a high SNR along with an excellent stability and size distribution for biological applications.

Clinical imaging probes that are biodegradable provide the significant advantage of being able to be broken down in the body and removed after they have served their function. To demonstrate that bioharmonophores are indeed biodegradable, we utilized the highly effective serine protease, proteinase K, which exhibits a broad cleavage specificity^18^. We incubated bioharmonophores with a proteinase K concentration that is routinely used for dissolving tissue structures^19^ and probed the extent of degradation by monitoring the SHG signal at different time intervals (**Fig. 3a**). We observed a decrease of SHG signal within 2 hours of protease incubation. After 10 hours, the SHG signal disappeared and the turbid bioharmonophore suspension became transparent (**Fig. 3b and Supplementary Fig. 7**), indicating a successful biodegradation of the bioharmonophore.

**Figure 3.**
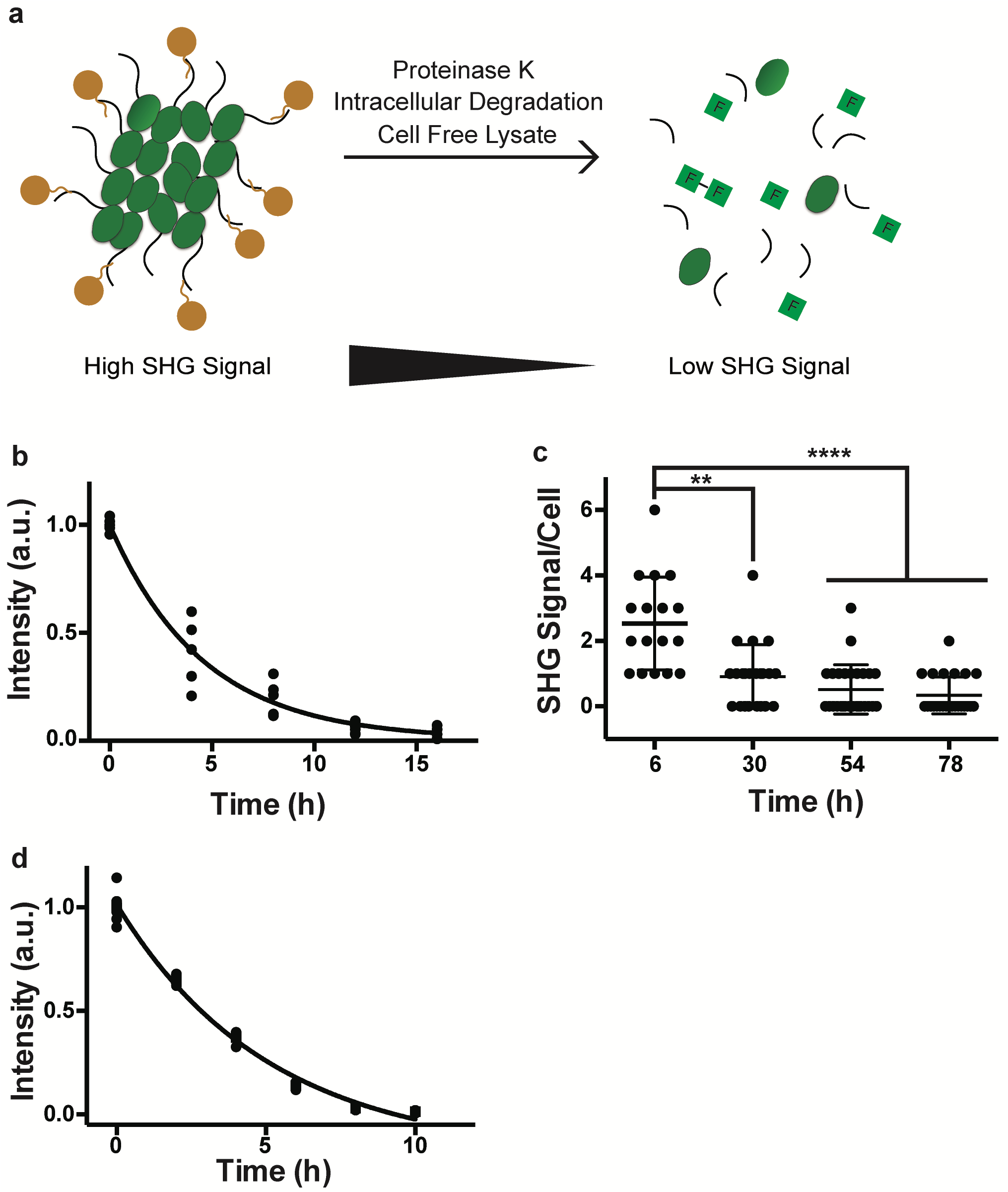
Bioharmonophores can be degraded by proteases, cells, and cell-free lysate systems. **a**, Schematic showing different degradation methods utilized to assess biodegradability of the bioharmonophores. **b**, Graph displaying the change of SHG signal intensity over time of bioharmonophores incubating with proteinase K (n=5). Mean values of data points were fitted for one phase exponential decay. **c**, Quantification of SHG signal/cell after overnight incubation of Tat-peptide functionalized bioharmonophores over time. SHG signal/cell is significantly reduced 30 hours after reseeding. Mean ± s.d. ****, P < 0.0001, **, P < 0.005, *, P < 0.05 (non-parametric Kruskal-Wallis test with Dunn’s post hoc multiple comparison). **d,** Graph showing the loss of SHG signal intensity when bioharmonophores are subjected to the cell-free reticulate lysate degradation system (n=5). Mean values of data points were fitted using a one phase exponential decay. Scale bar, 10 μm (**c**); Scale bar, 10 μm (**d**).

To evaluate bioharmonophore degradation under physiological conditions (**Fig. 3c**), we functionalized bioharmonophores with Tat-derived cell penetrating peptides^20^ using bioorthogonal click chemistry, (**Supplementary Fig. 8**), and incubated them with a model cancer cell line (see below) overnight. Adherent cells were then detached by trypsinization, and centrifuged to remove excess bioharmonophores that did not enter the cancer cells. Following this procedure, cells were reseeded and fixed at specific time periods to monitor bioharmonophores degradation (i.e. the intracellular presence of SHG signal per cell) using nonlinear optical imaging. 30 hours after cell reseeding, a pronounced decrease of intracellular SHG signal per cell was noticeable (**Fig. 3c**). As bioharmonophores displayed long-term photostability even at low pH values (**Supplementary Fig. 9**), the drop of signal was not due to their potential accumulation in acidic endolysosomal compartments over time. In order to show that bioharmonophores can be degraded using intracellular proteolytic degradation, we tested whether the bioharmonophores could be degraded using a cell-free lysate system based on an established cell free degradation assays^21^. We also observed reduced SHG signal, indicating that intracellular enzymatic degradation of bioharmonophores might account for the signal loss (**Fig. 3e and Supplementary Fig. 10**). Importantly, bioharmonophores did not exhibit any short-term toxicity *in vitro* and *in vivo* (**Supplementary Fig. 11**) and did not induce protein aggregation^22^ (**Supplementary Fig. 12**), rendering them safe imaging probes.

Among various diagnostic applications, bioharmonophores could be ideal imaging probes for single-cell cancer detection due to their high SNR and photostability, which other intravital imaging modalities cannot achieve^3^. To demonstrate the unique optical features of bioharmonophores for cancer targeting and imaging, we employed xenograft zebrafish cancer models, which offer speed, cellular resolution, and the ability to perform large numbers of transplants for obtaining valuable information about several cancer types^23, 24^.

To generate a highly aggressive cancer model that can be tracked over time, we injected a DsRed-expressing metastatic human melanoma cells (MDA-MB-435-DsRed) into the Duct of Cuvier (DoC) of zebrafish embryos at 2 dpf (days post fertilization)^23, 24^ (**Fig. 4a**). By 3 days after the injection, the resulting tumors spread to various locations in the body and were found next to blood vessels, which likely support the tumors with nutrients^25^ (**Fig. 4b**).

**Figure 4.**
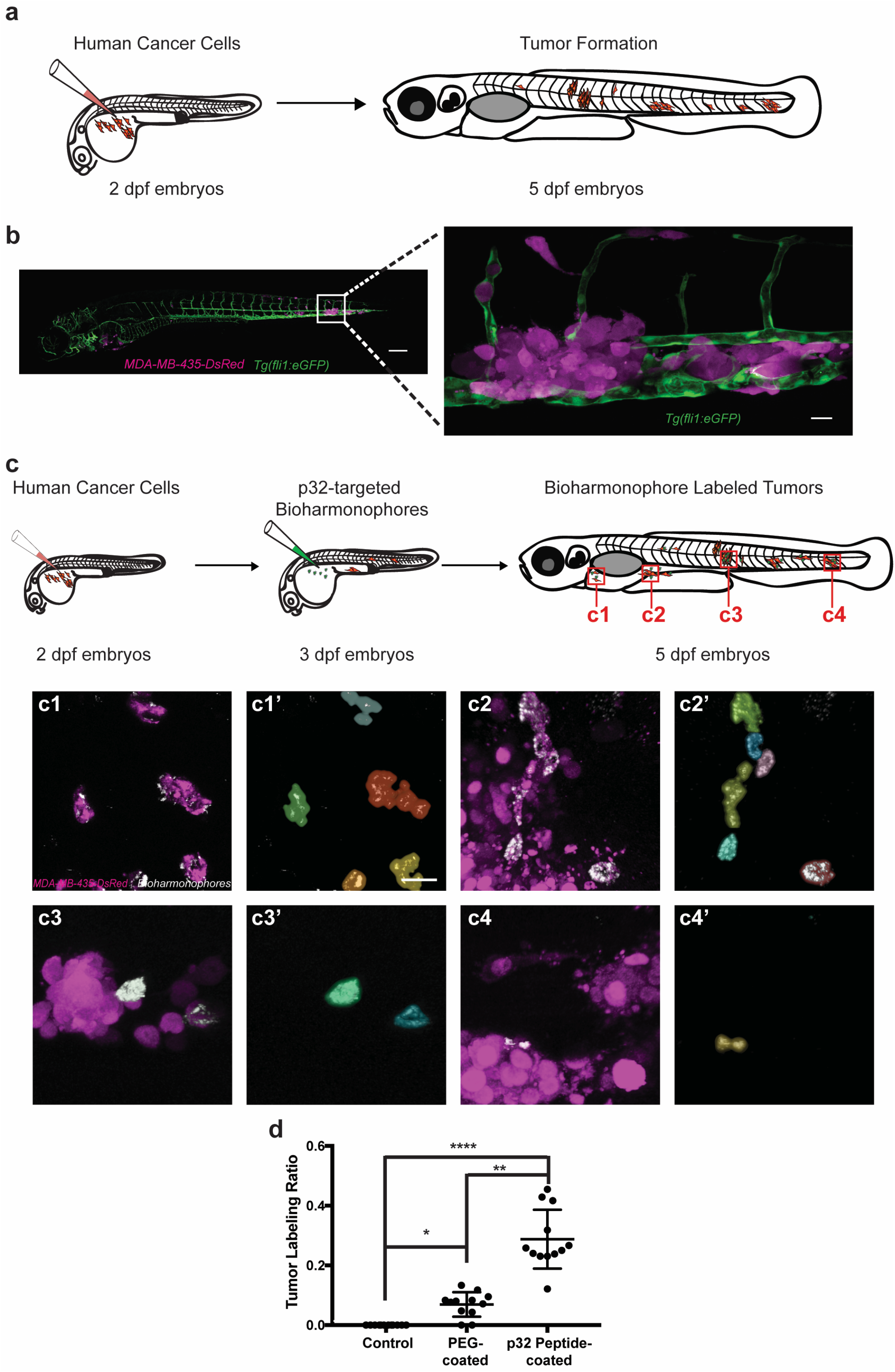
Bioharmonophores can be specifically targeted to single cancer cells *in vivo*. **a**, Schematic showing the generation of a zebrafish cancer model by injecting MDA-MB-435-DsRed cancer cells into the Duct of Cuvier (DoC) at 2 dpf, resulting in tumors spread to multiple locations of the zebrafish body at 5 dpf. **b**, Composite image of the cancer model (left) in a 5 dpf old zebrafish embryo. Close-up image of one of the tumor sites (right) reveals DsRed-labeled tumors (magenta), adjacent to the eGFP-labeled vasculature (green). **c**, Schematic showing cancer cell injection of 2 dpf zebrafish embryos followed by bioharmonophore injection into DoC of 3 dpf zebrafish embryos and subsequent fluorescence and SHG imaging at 5 dpf. Red rectangles labeled as **c1-4** denote the regions of interest that are illustrated in more detail. Individual panels showing the images of labeled cancer cells with the details of bioharmonophore (white) labeling down to single cancer cells (magenta) in solid tumors **(c1-4)**. Colored cell boundary reconstruction of targeted cancer cells using the bioharmonophore SHG signal **(c1’-4’)**. Note that cellular bioharmonophore distribution can in most cases predict cell morphologies. Scale bar, left panel 200 μm, right panel 20 μm (**b**); Scale bar, 15 μm (**c**). **d,** Quantification of the fraction of SHG-labeled tumors as the ratio of labeled tumors to all tumors in a given zebrafish embryo after PEG- and p32 peptide-coated bioharmonophore injection, respectively. Each data point signifies one zebrafish. Note that active targeting with p32-coated bioharmonophores significantly increases the labelling efficiency (approx. 4-fold). Mean ± s.d. ****, P < 0.0001, **, P = 0.0063, *, P = 0.0470 (non-parametric Kruskal-Wallis test with Dunn’s post hoc multiple comparison). N=12, pooled from 3 independent experiments.

To demonstrate the specificity and efficiency of bioharmonophores as novel contrast agents that can accomplish resolution down to the single cell *in vivo*, we targeted bioharmonophores to tumor sites by taking advantage of the surface protein p32/gC1qR as a unique molecular marker for MDA-MB-435-DsRed cells^26^. To this end, we functionalized bioharmonophores with a p32 targeting peptide, injected them into the DoC of zebrafish embryos at 3 dpf, one day after the embryos were injected with MDA-MB-435-DsRed cancer cells, (**Fig. 4c**) and assessed colocalization between cancer cells and bioharmonophore signal at 5 dpf (see **Supplementary Note 2** - Determining the fraction of bioharmonophore-labeled tumors in a zebrafish cancer model). In the absence of tumors, functionalized bioharmonophores did not cause clustering at the site of injection and were localized at different parts of injected zebrafish embryos (**Supplementary Fig. 13d-i**), indicating good biodistribution. Without bioharmonophore injection, zebrafish as well as tumor sites did not produce any SHG background signal (**Supplementary Fig. 13a-c, 14a-c**) with the exception of minimal endogenous SHG signal localized at the zebrafish tail^27^, which was excluded from assessing specificity of tumor targeting (**Supplementary Fig. 15**). In the case of passive targeting, zebrafish injected with PEG-coated bioharmonophores revealed limited tumor labeling (**Supplementary Fig. 14d-f**), stemming from leaky blood vessels and enhanced permeability and retention effect (EPR)^28^. In contrast, we observed an increased accumulation of p32 peptide-targeted bioharmonophores within individual cancer cells at tumor sites throughout the zebrafish embryos (**Fig. 4c1-c4’**), indicating that the tumor-labeling specificity and efficiency is highly dependent on the p32 targeting peptide. While p32 peptide-targeted bioharmonophores can extravasate to different tumor sites, not all the cancer cells were labeled (**Fig. 4c**). This observation is potentially due to limited accessibility within densely packed solid tumors^29^ and the continued proliferation and metastasis of cancer cells between bioharmonophore administration and imaging (see **Supplementary Note 2** - Determining the fraction of bioharmonophore-labeled tumors in a zebrafish cancer model).

In order to determine the extent of labeling of targeted bioharmonophores in the xenograft zebrafish cancer model, we measured the colocalization of cancer cells with bioharmonophores at each tumor site for non-injected zebrafish as well as for zebrafish that were injected with p32 peptide-targeted and PEG-coated bioharmonophores, respectively (see **Supplementary Note 2** - Determining the fraction of bioharmonophore-labeled tumors in a zebrafish cancer model, **Fig. 4c,d**). The number of tumors were not significantly different between datasets (**Supplementary Fig. 16**). The zebrafish cancer model injected with p32 peptide-targeted bioharmonophores had a significantly higher fraction of labeled tumors compared with non-injected and PEG-coated bioharmonophores (**Fig. 4d**) due to our active targeting strategy. Overall, these results demonstrate that bioharmonophores exhibit high SNR and outstanding photostability for efficient labeling of individual cancer cells at multiple tumor sites *in vivo*.

In summary, we introduced bioharmonophores as a novel class of imaging probes that retain all the photophysical advantages of previously introduced inorganic SHG nanoprobes. Because bioharmonophores consist of a biodegradable peptide core and a polymer shell, they can be metabolized within cells, which could render them the ideal contrast agent for clinical imaging applications. The straightforward implementation of robust functionalization strategies and a sufficiently high metabolic stability *in vivo* allowed us to target bioharmonophores with high detection sensitivity to individual tumor cells in live zebrafish embryos. With the recent development of nonlinear microendoscopes^30, 31^, bioharmonophores have the potential to emerge as superior contrast agents during image-guided surgery to help surgeons perform safer and highly precise tumor removal procedures. Moreover, their unique ability to target single cells could be exploited for detecting cancer stem cells, a subpopulation of cells responsible for tumorigenicity, invasion, and metastasis^32^. Once successfully identified, the nonlinear signal of bioharmonophores could be used for light induced drug delivery or photodynamic therapy^33^. By employing pulsed lasers in the infrared wavelength range that permit deep tissue penetration, targeted bioharmonophore signal could trigger highly localized cancer stem cell death^34^.

Finally, as the SHG signal intensity of bioharmonophore relies on the unique self-assembly behavior of each peptide^35^, we anticipate that a screen for alternative peptide sequences may yield even brighter bioharmonophores that will potentially permit diagnosis with deep-tissue single-molecule detection sensitivity.

## Acknowledgements

We thank members of the Pantazis group for discussion and feedback. We thank W.P. Dempsey for feedback on the manuscript. We thank the Scientific Center for Optical and Electron Microscopy (ScopeM) for their help in imaging bioharmonophores. We also thank T. Weber of the Crystallography Laboratory of ETH Zurich for his help with the XRD analysis. We thank R. Klemke for kindly providing the MDA-MB-435-DsRed cell line. We thank M. Affolter and H.G. Belting for providing some of the zebrafish eggs. This work was supported by the Swiss National Science Foundation (SNF grant no. 31003A_144048), the European Union Seventh Framework Program (Marie Curie Career Integration Grant (CIG) no. 334552), and the Swiss National Center of Competence in Research (NCCR) “Nanoscale Science”, which were awarded to P. Pantazis who is also a Royal Society Wolfson Research Merit Award holder.

## Author contribution

A.Y.S. conceived and A.Y.S. and P.P refined the idea. A.Y.S. produced and characterized bioharmonophores with the help of S.J., S.B., D.C., and K.L.. A.Y.S., C.T., and S.R., performed optical characterization. S.Y. and A.Y.S. performed cell culture experiments. A.Y.S. and S.Y. generated *in vitro* and *in vivo* imaging data. M.K. and S.S. generated the zebrafish cancer model, performed bioharmonophore injections with the help of C.L.. A.Y.S. and P.P. wrote the manuscript and all authors contributed to editing the manuscript. P.P. supervised the project.

## Methods

### Formation of large-scale peptide nanotubes

Diphenylalanine (FF) and triphenylalanine (FFF) (Bachem) peptide assemblies were prepared as previously described^36^. Briefly, peptides were freshly dissolved in hexafluoroisopropanol (Sigma) at 100 mg/ml concentration prior to experiments and diluted to 5 mg/ml final concentration in deionized water.

### Encapsulation of SHG active peptide assemblies

For the evaluation of different peptides and their SHG capabilities, 30 mg PLLA was dissolved in 3 ml chloroform (Sigma) along with 15 mg triphenylalanine, 30 mg pentalalanine (Bachem), and 30 mg trileucine (Sigma) peptides in separate glass vials. Resulting suspension was mixed with aqueous SDS (Sigma) solution with a final 0.3% SDS concentration. A macroemulsion was obtained by stirring the samples at 1000 rpm for 1 hour. Afterwards, the samples were sonicated (Branson Sonifier) with a 1,5 inch tip at 70% power in a pulsed mode (30 seconds ON and 10 seconds OFF) under ice cooling. The chloroform was evaporated from the obtained emulsions by stirring the samples at 500 rpm at 40 °C overnight. For the remaining experiments with triphenylalanine peptide containing bioharmonophores, the same protocol was followed unless stated otherwise. For probing the optimal conditions for nanoparticle formation, FFF peptide, PLLA, and SDS concentrations were varied as described in Supplementary Figures 2, 3, and 4.

### Characterization of encapsulated SHG active peptide assemblies

Produced samples were characterized using Dynamic Light Scattering. Nanoparticle morphology, aggregation tendency along with the SHG signal intensity were evaluated using nonlinear microscopy. XRD patterns were obtained using a PANalytical X’PERT Pro powder diffractometer in Bragg-Brentano geometry and with Cu K-alpha1 radiation in grazing incidence geometry between 2–60 using a step size of 0.0167. The samples were air-dried on silicon single crystals and four identical scans are obtained from each sample and summed up.

### SHG polarimetry

The SHG polarimetry was performed on a wide-field SHG microscope (See Supplementary Info). A 1030nm laser, pulse width 190fs, and 200kHz repetition rate (Pharos, LightConversion), delivered 36mW on the sample over a 150um FWHM diameter field-of-view (1 mJ.cm^−2^). Two noncolinear beams are incident on the sample, with an angle 30 degrees in between the two. SHG signal was detected in the phase matching condition (transmission). The image was recorded with an electron-multiplying intensified charge-coupled device (EM-ICCD) camera. Nonlinear polarimetry was performed by controlling and analyzing the polarization state of the illuminating and emitted beams. A polarization state generator, comprising a half- and a quarter-wave plate, was used. The polarization state of the emitted light was analyzed with a half-wave plate placed in the emission path, followed by a polarizing beam splitter.

### Second Harmonic Spectroscopy Patterns

SHG emission pattern measurement was performed on a custom-build setup for this purpose (See Supplementary Info). Excitation was performed with a 1030nm laser, pulse width 190fs, and 200kHz repetition rate, which delivered 60mW on the sample, a cylindrical cuvette containing the solution, over a 36um focal spot (30 mJ.cm^−2^). The signal was detected with a rotating PMT and a filter (515+10, Chroma) at angles between −90 and 90. Both incident and detection polarizations can be controlled.

### Stability of biodegradable bioharmonophores at different pH values

To evaluate how different pH values might influence the PLLA coated peptide assemblies and their signal intensity, synthesized bioharmonophores were centrifuged for 3 minutes at 13500 rpm and resuspended in citric acid/Na_2_HPO_4_ buffer ranging from 4 to 7 pH values. The bioharmonophores were incubated for 72 hours in the buffers containing 1% Tween 80 to prevent aggregation and the signal intensity was monitored using nonlinear microscopy.

### Biodegradation of bioharmonophores *in vitro*

Bioharmonophores were centrifuged for 3 minutes at 13500 rpm and resuspended in 1% Tween 80 containing PBS. In order to assess proteinase K (Sigma) degradation, 1 ml bioharmonophore suspension was incubated with 100 μg/ml final proteinase K concentration at 37 °C and the SHG signal intensity was measured every 2 hours. Similarly, in order to assess how bioharmonophores were degraded using cellular content, an ex vivo biodegradation protocol was adapted based on the Rabbit Reticulocyte Lysate system (Promega). In a typical setup, 1 ml of bioharmonophore in 1% Tween 80 containing PBS was mixed with 25 mM phosphocreatine (Sigma), 10 μg/ml phosphocreatine kinase (Sigma), 1 mM ATP (Sigma), and 50 μl Rabbit Reticulocyte Lysate. The mixture was incubated at 37 °C and the SHG signal intensity was monitored every 2 hours.

### Biodegradable bioharmonophore functionalization

1 ml of 1.5 mg/ml bioharmonophores were incubated with 1 mg Candida Antarctica Lipase B (Sigma) for 2 hours, which hydrolyzes the PLLA polymer to increase the number of carboxyl groups. Bioharmonophore suspension was centrifuged at 13500 rpm for 3 minutes and resuspended in 1% Tween 80 containing PBS, and mixed with 10 mg *N*-(3-dimethylaminopropyl)-*N*′-ethylcarbodiimide hydrochloride (EDC) (Sigma), 10 mg *N*-hydroxysuccinimide (NHS) (Sigma), and 10 mg methoxypolyethylene glycol amine 5000 Da (mPEG Amine) (Sigma) for 2 hours. The suspension was centrifuged and resuspended in 1% Tween 80 containing PBS and stored at 4 °C prior to use.

For further functionalization experiments thiol-PEG-amine 2000 Da (SH-PEG-NH_2_) (Sigma) was used as a platform for bioorthogonal click chemistry. In a similar setup, 10 mg EDC, 10 mg NHS, and 10 mg thiol-PEG-NH_2_ were incubated for 2 hours. The suspension was centrifuged and resuspended in 1% Tween 80 containing PBS with methyltetrazine-PEG4-Maleimide (Click Chemistry Tools) of 200 μM final concentration. The mixture was incubated for 2 hours at room temperature, centrifuged, and resuspended in 1% Tween 80 containing PBS.

The other click chemistry pair trans-cyclooctene (TCO)-PEG3-Maleimide (Click Chemistry Tools) (3 mM in 200 μl PBS) was incubated for 2 hours with cysteine-containing Tat or P32 targeting peptides (1 mM final concentration) depending on the application. The peptides were passed through Illustra Microspin G25 columns (GE Healthcare) to remove TCO-PEG3-maleimide.

200 μl tetrazine modified bioharmonophores were incubated with 20 μl TCO modified peptide for 2 hours. The bioharmonophore suspension was washed with 1% Tween 80 containing PBS to remove unbound peptides and resuspended in PBS to be immediately used for cell culture experiments.

### Cellular degradation and toxicity

MDA-MB-435-DsRed cancer cells were kindly gifted by Prof. R. Klemke. The cells were cultured at 37°C, 5% CO_2_, in high glucose DMEM with GlutaMAX (10569010, Thermo Fisher), supplemented with 10% FBS (P40-37500, Pan Biotech) and 1X Penicillin-Streptomycin solution (15140122, Thermo Fisher).). The cells were cultured on 6-well plates (140675, Thermo Fisher) until they reached ~80% confluency and were incubated with 400 μl Tat-derived cell penetrating peptide coated bioharmonophores overnight. The cells were washed with 1X PBS twice and detached using 0.05% Trypsin-EDTA (25300054, Thermo Fisher) in order to remove bioharmonophores that did not enter the cancer cells. Detached cells were centrifuged for 5 minutes at 500xg to remove excess bioharmonophores that were not taken up, reseeded or ibitreat coated 8-well slides (80826, Ibidi GmbH), and fixed after 6, 30, 54 and 78 hours to monitor bioharmonophores degradation. The samples were then washed 3 times with 1X PBS and stained with the CellMask Orange Membrane dye (Invitrogen). The samples were washed again and imaged subsequently. To determine cell viability after treatment with functionalized bioharmonophores, trypan blue exclusion method was used. Briefly, cells in triplicates seeded in 96-well tissue culture plates (167008, Thermo Fisher) were exposed to varying concentrations of functionalized bioharmonophores for 48 or 72 hours. After incubation, cells were washed with 1X PBS twice and detached as described above. 10 μl of cell suspension was then mixed with 10 μl 0.4% Trypan Blue, and 4 μl of this mixture was added to the cell counting slide (C10228, Thermo Fisher) and measured using Countess II Automated cell counter (Thermo Fisher). The viability was expressed as a fold difference of the untreated samples for each time point.

### Toxicity Assay and Thioflavin T staining

For toxicity assay, cells were grown in 96 well plates and were incubated with bioharmonophores at different concentrations for 48 and 72 hours. After the incubation period, the cells were detached with trypsinization and their viability was analyzed using Trypan Blue (Sigma) staining.

For Thioflavin Staining cells were seeded in an 8-well chamber (ibidi) with 50% confluency. The cells were treated with either Amyloid Beta Peptide (Bachem) or 5 μl of bioharmonophores for 24 hours and extensively washed with PBS to remove excess peptides and bioharmonophores. To evaluate whether bioharmonophores induce fibril formation the cells were fixed with 4% paraformaldehyde for 10 minutes and washed with PBS three times. Afterwards, 0.05% Thioflavin T (Sigma) solution was added to the sample for 8 minutes and excess dye was washed with 80% ethanol for 5 minutes. The washing step was repeated three times and the samples were imaged using confocal microscopy.

### Zebrafish Cancer Model and bioharmonophore Targeting

Animal experiments and zebrafish husbandry were approved by the “Kantonales Veterinaeramt Basel-Stadt”. MDA-MB-435-DsRed cancer cells were injected into the Duct of Cuvier of *Tg(fli1:egfp)* zebrafish embryos at 2 days post fertilization (dpf). After injection, embryos were incubated for 1 hour at 29°C for recovery and cell transfer then verified by fluorescence microscopy. Fish harboring red cells were incubated at 35°C essentially as described before^23, 24^. Fish were anesthesized and embedded in low melting agarose as described previously^37^ and were imaged at 5 dpf for assessing cancer cell localization.

For targeting experiments, p32/gC1qR ligand-functionalized bioharmonophores were injected into the zebrafish embryos 24 hours after cancer cell injection following the same procedure. *In vivo* bioharmonophores targeting was evaluated at 5 dpf using nonlinear laser scanning microscopy.

### Transmission Electron Microscopy

Bioharmonophore samples were spun down to remove aggregated nanoparticles at 3000 rpm for 3 minutes and the bioharmonophores (i.e., the supernatant of the centrifuged solution). 5 μl of the sample was placed on a carbon coated grid (Quantifoil, D) previously glow-discharged for 30 seconds (Emitech K100X, GB). The drop was allowed to remain for 60 seconds; after this interval, excess fluid was drained along the periphery using a piece of filter paper followed by staining with 2% uranyl acetate for 1 second and 15 seconds, respectively. Excess moisture was drained after each step and when dry the grid was examined in an FEI Morgagni 268 TEM operated at 100 kV.

### Nonlinear and Confocal Light Microscopy

Bioharmonophores were immobilized in low melting agarose by mixing 200 μL bioharmonophore with 100 μL 1% SeaPlaque low melting agarose (Lonza) solution in 8-well imaging chambers (Lab-Tek). Imaging experiments were performed on a Zeiss LSM 780 microscope (Carl Zeiss AG) equipped with a spectral GaAsP detector and a tunable two-photon laser source (Chameleon Ultra II, Coherent Inc.), using an LD C-Apochromat 40x/1.1 water immersion objective lens (Carl Zeiss AG). Throughout the imaging experiments, bioharmonophores were illuminated with 850 nm incident wavelength and the SHG signal was collected between 405 and 435 nm or with a GaAsP spectral wavelength detector for spectral measurements.

### Statistical analysis

All numerical values represent mean ± s.d. Sample sizes (n) were given in the figure legends for each experiment. Each experiment was repeated at least 3 times. Normal distribution of datasets were established using D’Agostino & Pearson omnibus normality test where P>0.05 indicated Gaussian distribution. When all the datasets had Gaussian distribution, one-way Anova was used for multiple comparisons followed by Tukey’s multiple comparisons. When one or more datasets showed a non-Gaussian distribution or high degree of variance as in the case of zebrafish tumor models, Kruskal-Wallis test was applied along with Dunn’s multiple comparisons. For all statistical tests, P value was reported, n.s., P > 0.05, *, P < 0.05, **, P < 0.005, ***, *P*<0.001, ****, P < 0.0001. Second order polynomial fit and one phase exponential decay values were calculated and graphs were drawn using GraphPad Prism 6.

## References

1. Luker GD, Luker KE. Optical Imaging: Current Applications and Future Directions. Journal of Nuclear Medicine 2007, 49(1): 1–4.

2. Pantazis P, Maloney J, Wu D, Fraser SE. Second harmonic generating (SHG) nanoprobes for in vivo imaging. Proceedings of the National Academy of Sciences 2010, 107(33): 14535–14540.

3. Lindner JR, Link J. Molecular Imaging in Drug Discovery and Development. Circulation: Cardiovascular Imaging 2018, 11(2).

4. Koch M, Ntziachristos V. Advancing Surgical Vision with Fluorescence Imaging. Annual Review of Medicine 2016, 67(1): 153–164.

5. Lamberts LE, Koch M, de Jong JS, Adams ALL, Glatz J, Kranendonk MEG, et al. Tumor-Specific Uptake of Fluorescent Bevacizumab– IRDye800CW Microdosing in Patients with Primary Breast Cancer: A Phase I Feasibility Study. Clinical Cancer Research 2017, 23(11): 2730–2741.

6. Billinton N, Knight AW. Seeing the Wood through the Trees: A Review of Techniques for Distinguishing Green Fluorescent Protein from Endogenous Autofluorescence. Analytical Biochemistry 2001, 291(2): 175–197.

7. Dempsey WP, Fraser SE, Pantazis P. SHG nanoprobes: Advancing harmonic imaging in biology. BioEssays 2012, 34(5): 351–360.

8. Viskota JCuc, Dempsey WP, Fraser SE, Pantazis P. Surface functionalization of barium titanate SHG nanoprobes for in vivo imaging in zebrafish. Nature Protocols 2012, 7(9): 1618–1633.

9. Sugiyama N, Sonay AY, Tussiwand R, Cohen BE, Pantazis P. Effective Labeling of Primary Somatic Stem Cells with BaTiO 3Nanocrystals for Second Harmonic Generation Imaging. Small 2018, 14(8): 1703386.

10. Lakshmanan A, Zhang S, Hauser CAE. Short self-assembling peptides as building blocks for modern nanodevices. Trends in Biotechnology 2012, 30(3): 155–165.

11. Kholkin A, Amdursky N, Bdikin I, Gazit E, Rosenman G. Strong Piezoelectricity in Bioinspired Peptide Nanotubes. ACS Nano 2010, 4(2): 610–614.

12. Handelman A, Beker P, Amdursky N, Rosenman G. Physics and engineering of peptide supramolecular nanostructures. Physical Chemistry Chemical Physics 2012, 14(18): 6391–6408.

13. Staff RH, Schaeffel D, Turshatov A, Donadio D, Butt H-J, Landfester K, et al. Particle Formation in the Emulsion-Solvent Evaporation Process. Small 2013, 9(20): 3514–3522.

14. Rabotyagova OS, Cebe P, Kaplan DL. Role of Polyalanine Domains in β-Sheet Formation in Spider Silk Block Copolymers. Macromolecular Bioscience 2010, 10(1): 49–59.

15. Handelman A, Kuritz N, Natan A, Rosenman G. Reconstructive Phase Transition in Ultrashort Peptide Nanostructures and Induced Visible Photoluminescence. Langmuir 2016, 32(12): 2847–2862.

16. Handelman A, Lavrov S, Kudryavtsev A, Khatchatouriants A, Rosenberg Y, Mishina E, et al. Nonlinear Optical Bioinspired Peptide Nanostructures. Advanced Optical Materials 2013, 1(11): 875–884.

17. de Beer AGF, Roke S. Nonlinear Mie theory for second-harmonic and sum-frequency scattering. Physical Review B 2009, 79(15): 155420.

18. Tsuji H, Ogiwara M, Saha SK, Sakaki T. Enzymatic, Alkaline, and Autocatalytic Degradation of Poly(l-lactic acid): Effects of Biaxial Orientation. Biomacromolecules 2006, 7(1): 380–387.

19. Sepp R, Szabo I, Uda H, Sakamoto H. Rapid techniques for DNA extraction from routinely processed archival tissue for use in PCR. Journal of Clinical Pathology 1994, 47(4): 318–323.

20. Lewin M, Carlesso N, Tung C-H, Tang X-W, Cory D, Scadden DT, et al. Tat peptide-derivatized magnetic nanoparticles allow in vivo tracking and recovery of progenitor cells. Nature Biotechnology 2000, 18(4): 410–414.

21. Nguyen H, Gitig DM, Koff A. Cell-Free Degradation of p27 kip1, a G1 Cyclin-Dependent Kinase Inhibitor, Is Dependent on CDK2 Activity and the Proteasome. Molecular and Cellular Biology 1999, 19(2): 1190–1201.

22. Lee H-J, Shin SY, Choi C, Lee YH, Lee S-J. Formation and removal of alpha-synuclein aggregates in cells exposed to mitochondrial inhibitors. Journal of Biological Chemistry 2002, 277(7): 5411–5417.

23. Konantz M, Balci TB, Hartwig UF, Dellaire G, André MC, Berman JN, et al. Zebrafish xenografts as a tool for in vivo studies on human cancer. Annals of the New York Academy of Sciences 2012, 1266(1): 124–137.

24. Stoletov K, Kato H, Zardouzian E, Kelber J, Yang J, Shattil S, et al. Visualizing extravasation dynamics of metastatic tumor cells. Journal of Cell Science 2010, 123(13): 2332–2341.

25. Stoletov K, Montel V, Lester RD, Gonias SL, Klemke R. High-resolution imaging of the dynamic tumor cell–vascular interface in transparent zebrafish. Proceedings of the National Academy of Sciences 2007, 104(44): 17406–17411.

26. Agemy L, Kotamraju VR, Friedmann-Morvinski D, Sharma S, Sugahara KN, Ruoslahti E. Proapoptotic Peptide-Mediated Cancer Therapy Targeted to Cell Surface p32. Molecular Therapy 2013, 21(12): 2195–2204.

27. LeBert DC, Squirrell JM, Huttenlocher A, Eliceiri KW. Second harmonic generation microscopy in zebrafish. vol. 133. Elsevier, 2016, pp 55–68.

28. Nakamura Y, Mochida A, Choyke PL, Kobayashi H. Nanodrug Delivery: Is the Enhanced Permeability and Retention Effect Sufficient for Curing Cancer? Bioconjugate Chemistry 2016, 27(10): 2225–2238.

29. Jain RK, Stylianopoulos T. Delivering nanomedicine to solid tumors. Nature Reviews Clinical Oncology 2010, 7(11): 653–664.

30. König K, Ehlers A, Riemann I, Schenkl S, Bückle R, Kaatz M. Clinical two-photon microendoscopy. Microscopy Research and Technique 2007, 70(5): 398–402.

31. Sanchez GN, Sinha S, Liske H, Chen X, Nguyen V, Delp SL, et al. In Vivo Imaging of Human Sarcomere Twitch Dynamics in Individual Motor Units. Neuron 2015, 88(6): 1109–1120.

32. Yu Z, Pestell TG, Lisanti MP, Pestell RG. Cancer stem cells. Int J Biochem Cell Biol 2012, 44(12): 2144–2151.

33. Kachynski AV, Pliss A, Kuzmin AN, Ohulchanskyy TY, Baev A, Qu J, et al. Photodynamic therapy by in situ nonlinear photon conversion. Nature Photonics 2014, 8: 455.

34. Costa DF, Mendes LP, Torchilin VP. The effect of low- and high- penetration light on localized cancer therapy. Adv Drug Deliv Rev 2018.

35. Adler-Abramovich L, Gazit E. The physical properties of supramolecular peptide assemblies: from building block association to technological applications. Chem Soc Rev 2014, 43(20): 6881–6893.

36. Reches M, Gazit E. Controlled patterning of aligned self-assembled peptide nanotubes. Nature Nanotechnology 2006, 1(3): 195–200.

37. Dempsey WP, Fraser SE, Pantazis P. PhOTO Zebrafish: A Transgenic Resource for In Vivo Lineage Tracing during Development and Regeneration. PLOS ONE 2012, 7(3): e32888.

